# Workstation benchmark of Spark Capable Genome Analysis ToolKit 4 Variant Calling

**DOI:** 10.1101/2020.05.17.101105

**Authors:** Marcus H. Hansen, Anita T. Simonsen, Hans B. Ommen, Charlotte G. Nyvold

**Affiliations:** Haematolology-Pathology Research Laboratory, Research Unit for Haematology and Research Unit for Pathology, University of Southern Denmark and Odense University Hospital, Odense, Denmark; Department of Hematology, OUH, Denmark; Department of Hematology, Aarhus University Hospital (AUH), Denmark

**Keywords:** GATK4, Genome Analysis Toolkit, variant calling, next generation sequencing, Spark, Multithreading and parallel computing

## Abstract

**Background:** Rapid and practical DNA-sequencing processing has become essential for modern biomedical laboratories, especially in the field of cancer, pathology and genetics. While sequencing turn-over time has been, and still is, a bottleneck in research and diagnostics, the field of bioinformatics is moving at a rapid pace – both in terms of hardware and software development. Here, we benchmarked the local performance of three of the most important Spark-enabled Genome analysis toolkit 4 (GATK4) tools in a targeted sequencing workflow: Duplicate marking, base quality score recalibration (BQSR) and variant calling on targeted DNA sequencing using a modest hyperthreading 12-core single CPU and a high-speed PCI express solid-state drive.

**Results:** Compared to the previous GATK version the performance of Spark-enabled BQSR and HaplotypeCaller is shifted towards a more efficient usage of the available cores on CPU and outperforms the earlier GATK3.8 version with an order of magnitude reduction in processing time to analysis ready variants, whereas MarkDuplicateSpark was found to be thrice as fast. Furthermore, HaploTypeCallerSpark and BQSRPipelineSpark were significantly faster than the equivalent GATK4 standard tools with a combined ∼86% reduction in execution time, reaching a median rate of ten million processed bases per second, and duplicate marking was reduced ∼42%. The called variants were found to be in close agreement between the Spark and non-Spark versions, with an overall concordance of 98%. In this setup, the tools were also highly efficient when compared execution on a small 72 virtual CPU/18-node Google Cloud cluster.

**Conclusion:** In conclusion, GATK4 offers practical parallelization possibilities for DNA sequence processing, and the Spark-enabled tools optimize performance and utilization of local CPUs. Spark utilizing GATK variant calling is several times faster than previous GATK3.8 multithreading with the same multi-core, single CPU, configuration. The improved opportunities for parallel computations not only hold implications for high-performance cluster, but also for modest laboratory or research workstations for targeted sequencing analysis, such as exome, panel or amplicon sequencing.

## Introduction

The Genome Analysis Toolkit (1, 2) (GATK, Broad Institute, MA, USA) has been of tremendous value to bioinformaticians and biomedical research laboratories involved in next generation sequencing (NGS). The original GATK papers by McKenna et al. and DePristo et al. (1, 2) have been elaborated on and referenced extensively, emphasizing its wide usage in research. Not only did it provide a framework for consistent analysis of NGS data, but it also confronted the problem of inconsistent assignment of base quality scores by the different sequencing platforms. More importantly, GATK has also helped in the local realignment of reference genome mapped reads. It was observed that without these precautions, one in five called variants could be assigned false positives (2). From a clinical or a research point of view, the lack of such realignment is devastating for the downstream interpretation of somatic mutations in cancer detected with MuTect (3) or other somatic callers.

The preceding year has offered several advances to general bioinformatics and processing of sequencing data: First, 2018 offered new modestly priced chips, such as the 12 to 18-core CPUs launched in the fourth quarter by Intel and the 32-core by AMD. Secondly, solid state drives (SSD), which have been a necessity for practical handling of large sequencing files (4), have experienced a leap in read/write speed with the introduction of consumer disks utilizing PCIe 3.0 hardware interface. On the software side, the release of GATK4 (software.broadinstitute.org/gatk) with its fast Spark capabilities has coincided with several of these hardware innovations, while cloud computing has also become more powerful, flexible and easier to use for parallelization. Thus, multithreaded or parallel sequence processing on contemporary platforms mediates a potential leap in the amount of processed data per unit of time, when exploited optimally.

We benchmarked the quite novel parallelization of GATK4 based on Apache Spark (The Apache Software Foundation, MA, USA), which on one hand enables practical implementation in a high-performance cluster, and on the other, *it also enables efficient multithreading on single (or multiple) CPUs*, which is evaluated here. In theory, this eliminates the need for the scattering of the data into chunks or intervals in order to utilize the full potential of the available cores on the machine. In practice, this is still a reasonable approach to increase somatic calling speed with MuTect2 (3) in version 3.8 and GATK4. Albeit more efficient in the latter version, Mutect2 still awaits to be “*Sparkified”*. Behind the motivation and reasoning for local multithreading and parallelization lies the different needs in research, development and clinically applied bioinformatics, where a cluster may be impractical under some circumstances or may give rise to ethical concerns in clinical laboratory diagnostics. Thus, new possibilities to optimize GATK workflows on single machines are highly welcomed. For a recent benchmark of GATK4, not focusing on the Spark capabilities, see Heldenbrand et al. 2018 (5).

## Materials and methods

Data used in this benchmark was generated from 93 paired-end sequenced samples based on mononuclear cell DNA derived from patients diagnosed with acute leukemia. Targeted panel and amplicon sequencing were performed at Münchner Leukämielabor GmbH (Munich, DE) and Department of Molecular Medicine (Aarhus University Hospital, Aarhus, DK), using the NovaSeq and MiSeq platforms (Illumina, San Diego, CA, USA). The median read length was 140 and the median number of sequencing reads were 28.9×10^6^ (0.6×10^6^–117.1×10^6^). Paired-end whole exome sequencing data was downloaded from the 1000Genomes project data (6) (HG00121, read length: 90, unaligned and aligned reads: 52×10^6^). Compressed binary sequencing alignment files (BAM) were prepared by alignment to reference genome (GRCh37) using BWA (7) sorted by queryname or with subsequent coordinate sorting in Picard (Fig. 1).

**Figure 1.**
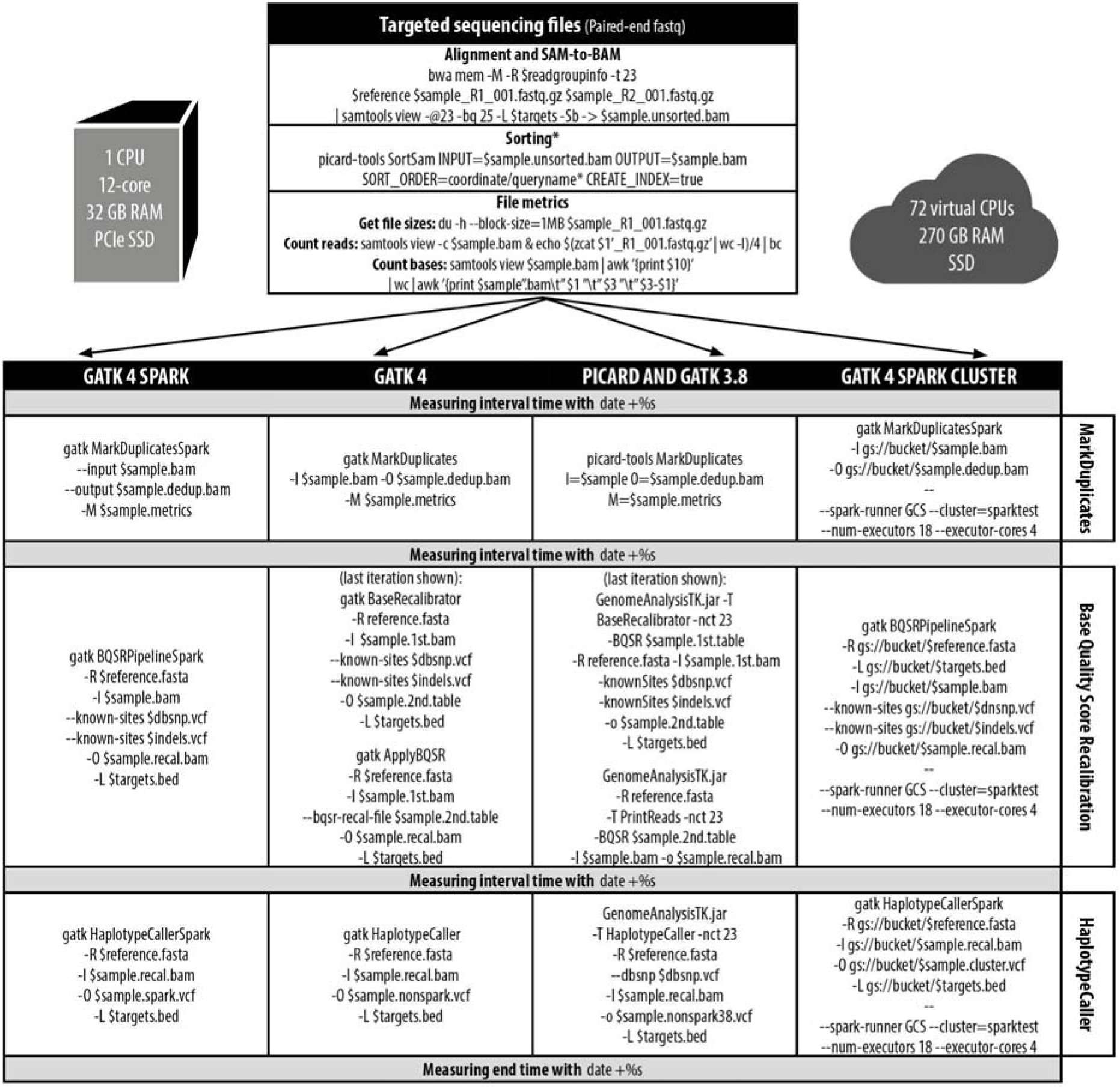
Benchmark workflow and commands. Coordinate sorted binary alignment files (BAM) were prepared from targeted paired-end sequencing and used in the benchmark test of variant calling workflow using GATK4 Spark, GATK4 and GATK3.8 with Picard MarkDuplicates 1.95. The local single CPU tests were conducted on a 12-core (24 logical cores) workstation with 32 GB RAM. In addition, a cloud cluster was configured with 72 virtual CPUs (18 nodes) to assess processing of whole exome sequencing data and the largest panel sequencing sample together with local workstation.

GATK4 Spark (version 4.0.12.0/4.1.1.0) on a stand-alone workstation was evaluated against the GATK4 non-Spark versions of MarkDuplicates, base quality score recalibration (BQSR) and HaplotypeCaller (n=93) in addition to GATK3.8 and Picard MarkDuplicates (version 1.95, default apt-get package) – the previous and thoroughly tested production tools used for targeted sequencing analyses on long term support Ubuntu version 14.04 at the hospital departments. GATK3.8 processing was stopped after an adequate number of the samples had been processed for comparison (n=59) due to exceedingly long execution times. In addition to panel sequencing, we evaluated the command-line tools on single exome sequencing samples using the current minor release (4.1.1.0). All the metrics were gathered locally on a clean installation of Ubuntu (14.04 LTS) using a single stand-alone mid-end workstation configured with a 12 core (24 logical cores) CPU (i9-7920X, 12×2.9GHz/4.3GHz boost, 16 MB cache, Intel, Santa Clara, CA, USA), 512 Samsung (Seoul, South Korea) GB PCIe SSD (up to 3500 MB/s read/2500 MB/s write) and 32 GB DDR4 quad-channel RAM. GATK4 commands were invoked through the supplied wrapper script, running Java SE 8 (JRE build 1.8, Oracle). A memory swap file of 64 GB was created on the PCIe SSD to tackle potential memory shortage. Disk performance and utilization during execution was additionally assessed with the linux *iostat* command.

Cloud evaluation of the Spark-enabled tools (version 4.1.1.0) was performed on a small virtual cluster with 18 nodes (Google Cloud (GC) Platform, Google LLC, Mountain View, CA, USA), running Ubuntu 18.04 LTS and Apache Spark 2.4, with 72 virtual CPUs (vCPUs) persistent SSD disks and 270 GB memory. Same GATK arguments as the local workflow described above were provided along with additional cluster specific arguments (Fig. 1). The number of executors and vCPUs, equivalent to the number of hyperthread cores, used for the comparison matched the number of nodes and available cores, respectively. The specific number of nodes used here was selected from performance testing of different node configurations with a fixed number of total vCPUs (i.e. 18×4, 9×8 and 4×18) and GATK4 Spark processing speed, hence the implementation of 72 vCPUs. Performance was evaluated based on the public exome data set (HG00121 (6)). In addition, downsampling (15–75%) of the largest aligned panel sequencing data file used in this benchmark (5.3 GB BAM) to assess the idle initiation times of MarkDuplicateSpark, BQSRPipelineSpark and HaplotypeCallerSpark on the cluster. Reference genome, list of targeted regions and data files were loaded from GC data storage buckets upon execution. All exome processing and cloud tests were performed in triplicates.

## Results

In this benchmark report we evaluate the multithreading capabilities of three of the most important tools in GATK4. Whereas the previous multithreading options in GAT3.8, not utilizing Spark, have largely been dropped, partly owing to its poor scalability and stability, the current version introduces sophisticated and unprecedented scaling possibilities, which is suited for contemporary hardware. This also holds potential for large scale analyses using cloud computing and clusters.

The benchmark results showed a decrease in processing time with both the non-Spark and Spark tools, and a linear relationship between the number of handled bases and the amount of time spent per sample. GATK4 MarkDuplicates was twice as fast as the early Picard MarkDuplicates (version 1.95) with a superior correlation between execution time and the number of bases processed (Fig. 2A), whereas MarkDuplicatesSpark from *coordinate* sorted alignments showed an overall poor correlation and decreased efficiency in this setup with roughly ten million bases processed per second, estimated from the median rate. Subsequently, we benchmarked alignments sorted according to *queryname* and the new documentation guidelines in version 4.1.1.0, which offered a stable version of MarkDuplicatesSpark. This provided a consistent and significant increase in speed (Fig. 2B, see Fig. 4A for exome data) by a factor of 1.7 (∼3.6×10^7^ b/s), compared to MarkDuplicates (∼2.1×10^7^ b/s)), and could utilize BWA output directly. Output from high-depth amplicon sequencing gave rise to a few outliers with lower speed.

**Figure 2.**
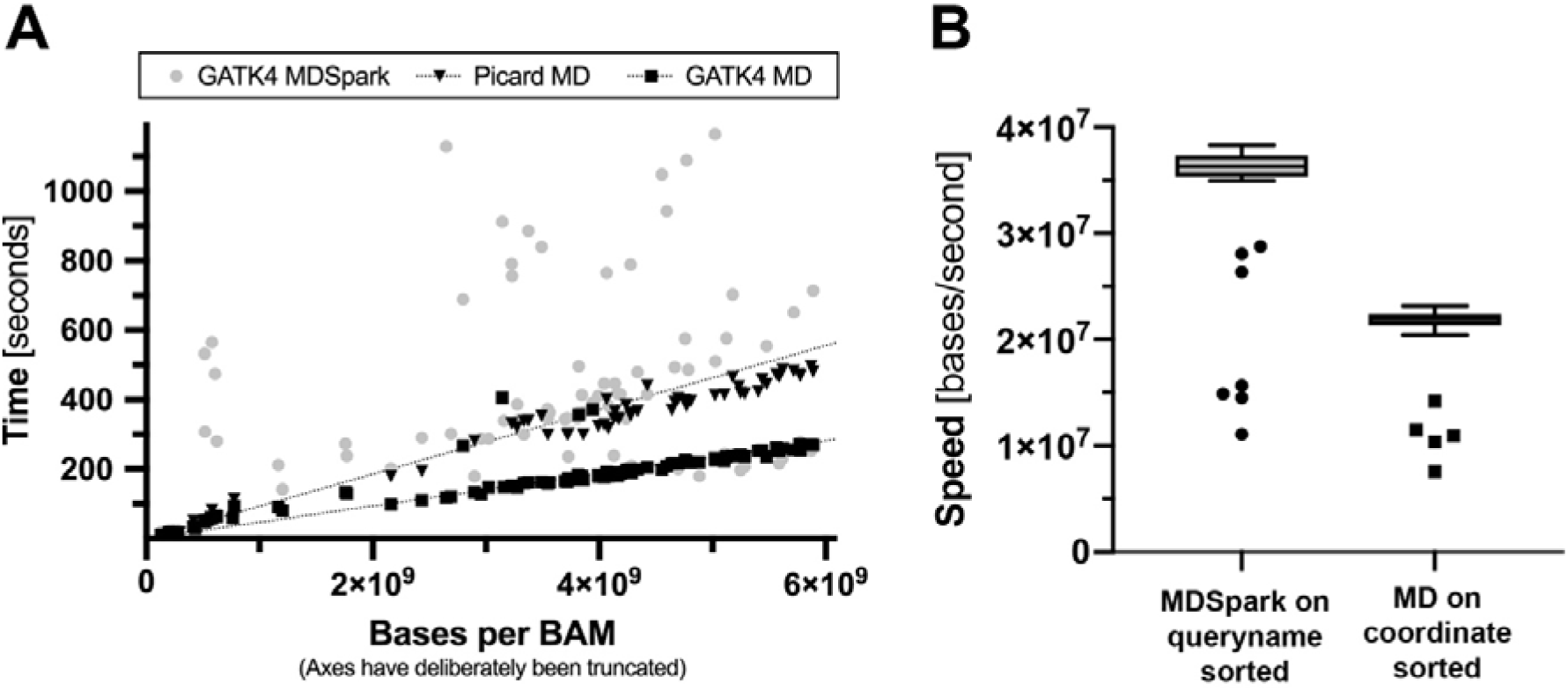
Performance of MarkDuplicates (MD). A doubling in processing speed was observed, when comparing GATK4 MarkDuplicates with Picard MD. Both tools displayed a linear correlation between elapsed time and the number of bases processed **(A)**, while the Spark version did not when using coordinate sorted alignments. The calculated rates were approximately 2.1×10^7^ bases per second (b/s) for GATK4 MD and 1.1×10^7^ b/s for Picard. The median rate for MarkDuplicatesSpark (MDSpark) on *groupname* sorted alignment was estimated to be 3.6×10^7^ b/s or 1.7 times faster than the single-threaded GATK4 MarkDuplicates **(B)**. The few outliers arise from amplicon sequencing.

**Figure 3.**
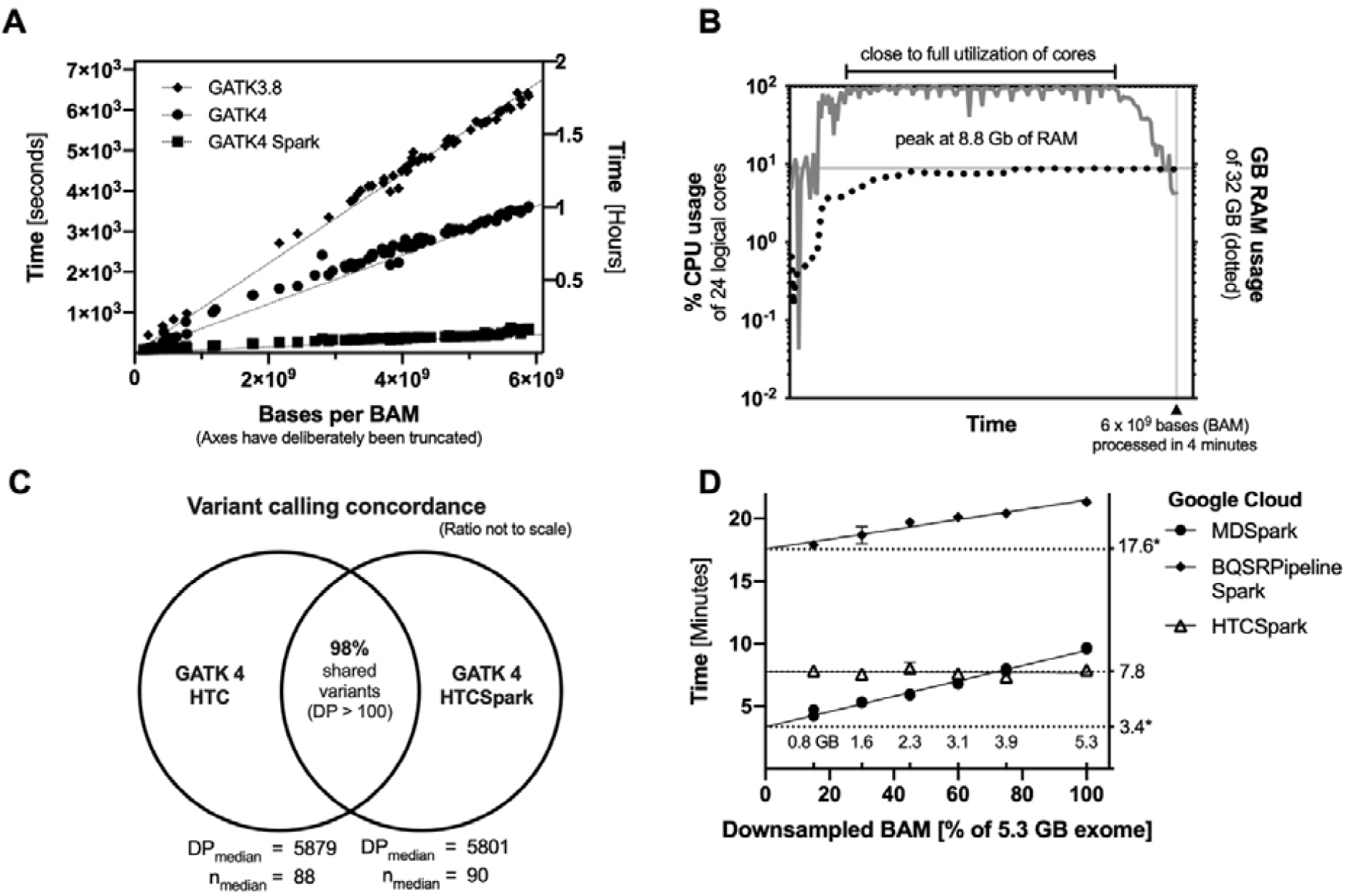
Benchmark of Spark variant calling and efficiency. GATK4 Spark quality score calibration and variant calling combined (BQSRPipelineSpark and HaplotypeCallerSpark) was found to be several times faster than the non-Spark and GATK3.8 counterparts **(A)**, with both tools efficiently utilizing multithreading and with low memory consumption. The CPU and memory usage of HaplotypeCallerSpark based on a random targeted sequencing sample is shown **(B)**. Read/write utilization (IO) of the PCIe solid state drive (up to 3500 MB/s read/2500 MB/s write) was found to be low (shown in dark gray) during execution of the GATK4 MarkDuplicateSpark (MD), base quality score recalibration (BQSRPipelineSpark) and HaplotypeCallerSpark (HTC), with the exception of a short 20 seconds burst during initiation of BQSR reaching approximately 100% disk utilization **(C**, median replicate values). The concordance of variant calling between Spark and non-Spark versions was found to be high **(D)**, with a similar number of variants detected (n_median_) and variant reads depths (DP_median_).

**Figure 4.**
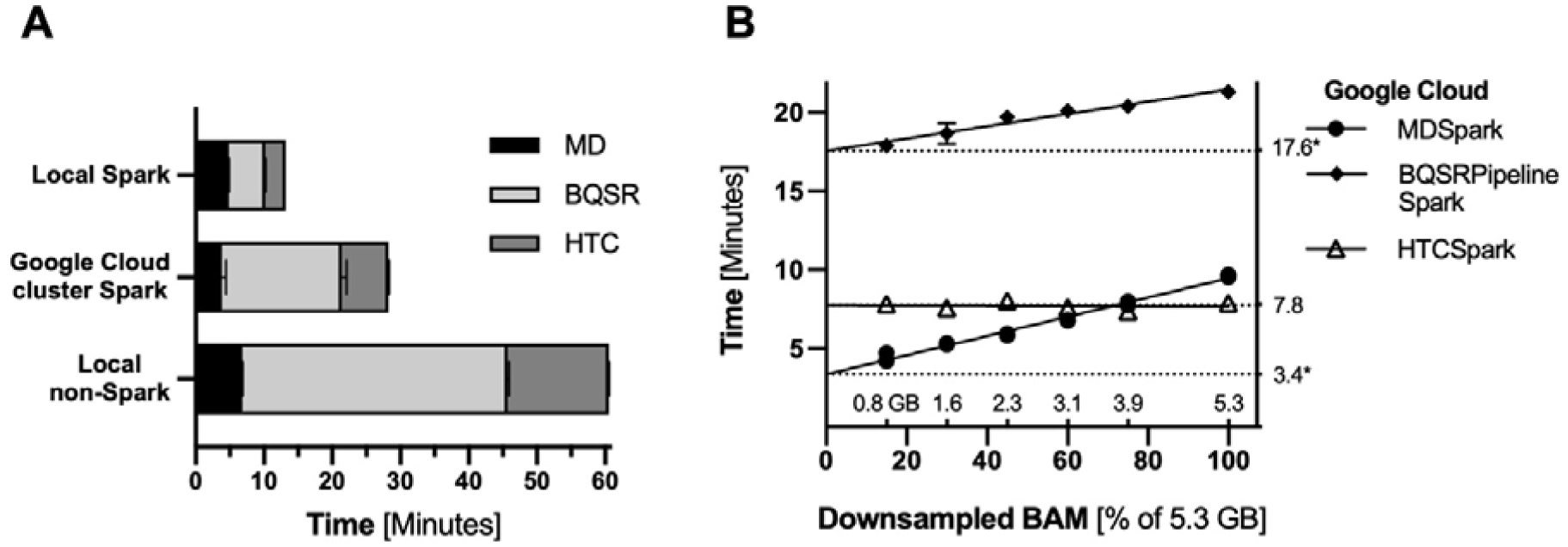
MarkDuplicatesSpark, BQSRPipelineSpark and HaploTypeCallerSpark were assessed using public exome data (HG00121), locally with a 12-core CPU and externally using a 72 virtual CPU/18-node cluster configuration (Google Cloud platform). Each analysis was performed in triplicates with high reproducibility. The local Spark execution time was approximately twice as fast as the cluster and four times as fast as the non-Spark GATK4 tools. We found that initiation of the cluster processing consumed a large part of the cluster wall-times (*) for MarkDuplicatesSpark and BQSRPipelineSpark. HaplotypeCallerSpark wall-time did not change with increasing number of reads from a 5.3 GB sequencing file.

Combined base quality score recalibration and variant calling for panel and amplicon sequencing were seven to eight (6.7–8.0) times faster than the non-spark GATK4 counterpart (Fig. 3A, see Fig. 4A for exome data), based on the median number of bases per second (b/s). Both tools were superior in terms of processed bases per unit of time, in comparison to GATK3.8, with nearly two (1.7–1.9) and up to 15-fold (11.15–14.6) increase in efficiency with the GATK4 non-Spark and Spark versions, respectively. The CPU usage in HaplotypeCallerSpark was close to full utilization of all 24 logical cores in major parts of the procedure (Figure 3B, randomly drawn sample) with less than 10 GB of RAM used at any time point in this setup. Likewise, BQSRPipelineSpark showed hefty bouts of CPU activity at near maximum utilization in its runtime as well as low RAM usage. The median processing rate of BQSRPipelineSpark and HaplotypeCallerSpark, combined, was ten million bases per second, highly comparable to the tested larger exome sequencing data, with each separate processing rate 2.1×10^7^ and 2.0×10^7^ b/s.

Disk utilization is a potential limiting factor in the overall processing speed. However, the PCIe SSD read/write burden was found to be low during major parts of the GATK4 workflow in contrast to CPU load, with the exception of BQSRPipelineSpark initiation (Fig. 3C, 1.9 GB BAM).

### Variant concordance between GATK versions

Next, we compared the panel sequencing variant output from GATK4 HaplotypeCaller with the respective variant call format (VCF) files generated by HaplotypeCallerSpark (Fig. 3D) and the 3.8 version of HaplotypeCaller. The median concordance, calculated as the fraction of GATK4 variants identical to the observed variants for each set, was in the first case found to be 0.98 (Q1=0.90, Q3=0.99) and 0.99 (0.96, 1) with or without indels, respectively, at a read depth threshold of 100 selected from a median depth 5801 (2629–6506). The median number of variants were 88 for GATK4 HaplotypeCaller and 90 for HaplotypeCallersSpark. The variant concordance to HaplotypeCaller 3.8 was found to be 0.94 (0.92, 0.97) and 0.96 (0.94, 0.98), with 89 variants called. In each case, the number of called variants compared to standard GATK4 HaplotypeCaller was not significantly different (p_Mann–Whitney_ > 0.05) and the concordance was found to be largely invariant to decreasing read depth thresholds.

### Implementing Spark workflow for exome sequencing and the Google Cloud Platform

Based on the results from the benchmark, we tested the Spark workflow on public whole exome sequencing data (HG00121) – from unaligned reads (52 million reads of 90 bases each) to analysis ready variants. The workflow combined BWA and the most current GATK4 MarkDuplicatesSpark, BQSRPipelineSpark and HaplotypeCallerSpark. Processing was completed in 28 minutes (measured in triplicates), of which BWA used 15 minutes. The combined GATK4 Spark rate corresponded to 5.9×10^6^ b/s, or approximately 4 million reads processed every minute, and a 78% reduction compared to equivalent GATK4 non-Spark (Fig. 4A). Likewise, a 53% reduction relative to the effective 72 vCPU cluster processing rate was observed. However, a second test using downsampling (15-100% of 106×10^6^ aligned reads, 5.3 GB) to test reproducibility showed that idle script initiation, such as copying of the files from the databucket, consumed a large part of the cluster wall-time (Fig. 4B). Therefore, for a fair comparison to the local machine utilizing a faster PCIe SSD, the benchmark baseline was *adjusted* accordingly. The equivalent speed was found comparable to the stand-alone workstation. We also observed that random downsampling did not alter the wall-time of HaplotypeCallerSpark, at least for the presented setup. This indicates that the number of variants influence the execution time, rather than the depth of coverage. Furthermore, it may cautiously indicate that a proper implementation of HaplotypeCallerSpark may provide extremely fast variant calling of large alignment files, such as whole genomes or exome sequencing with high depth of coverage.

## Discussion

We observed that HaplotypeCallerSpark and BQSRPipelineSpark were significantly faster than the equivalent GATK4 standard tools with a combined 85% reduction in execution time, and with a median rate of one million processed bases per second. The called variants were found to be in close agreement with the non-Spark version, with an overall concordance of 98%. The tools also displayed a higher performance locally in comparison to the 72 virtual CPU/18-node cloud cluster. GATK4 Spark implementation of quality score recalibration and variant calling was efficient and matched, or even outperformed, the configured cloud cluster in terms of processing time. This may only be true for smaller sequencing files of a few gigabytes or less. We argue that once the GATK4 Spark tools enter stable release, HaplotypeCallerSpark and BQSRPipelineSpark will be highly suitable for targeted sequencing and even for fast processing of medium coverage exome sequencing on modern CPUs and disks. With the most optimal usage of the GATK4 tools subsequent to alignment, it is feasible to finish analyzing a whole exome or extensive panel sequencing with high depth of coverage and paired-end reads in less than one hour on a contemporary workstation, while following a workflow similar to current *Best Practices* with regard to sequencing preprocessing and germline variant calling (software.broadinstitute.org/gatk/best-practices). The results show that processing of approximately 24–48 typical whole exomes of 50–100 million paired-end sequencing reads can be performed in a single day – and a larger amount of typical deep panel sequencing samples.

From the analyses we found both the Spark and non-Spark version of GATK4 MarkDuplicates to be fast and highly suitable for production use, while the BQSRPipelineSpark and HTCSpark provided efficient processing with minor deviation from non-spark variant calling. As GATK4 MarkDuplicatesSpark performance has been a subject of online debate, we emphasize that sorting by *queryname* is extremely important before submitting the alignment to tagging or removal of duplicate reads. Of note, MarkDuplicatesSpark is suited for processing of BWA output directly, which makes the processing speed achieved here even more remarkable, and thus not directly comparable to the non-Spark version.

As of early 2019, most of the GATK Spark tools are still offered in a beta version, and some tools, such as MuTect2, do not yet have inherent capabilities for parallelization. With the recent release of version 4.1, MarkDuplicatesSpark has moved beyond its beta version. Currently, pipeline tools other than the BQSRPipelineSpark have been released, such as ReadsPipelineSpark which accepts unaligned BAM and outputs analysis ready variants (VCF). From our evaluation the usage of this pipeline tool is confined to experimental testing as we have experienced several crashes just before the generation of the VCF file. We have also experienced close to physical RAM depletion at runtime. Thus, we recommend using more than 32 GB of RAM, as some Spark pipeline procedures may be memory-intensive.

In perspective, the current possibilities of installing high-end workstations with 32–36 physical cores, e.g. by using two Intel Xeon, X-series processors or even a single 32-core CPU with AMD RYZEN ThreadRipper (Santa Clara, CA, USA, launched in 2018), may render the need for cluster access superfluous in many research or clinical laboratories. Such impetus and innovations, as described, offer a viable and economical alternative to large high-performance clusters or to Illumina’s 2018 soft- and hardware acquisition, DRAGEN from Edico Genome, which boasts to process an entire genome (30x coverage) in just 25 minutes (Illumina). It had already proven its worth in 2015, when setting the world record in clinical whole genome analysis (8), and again in 2018. While exciting, the analysis platform may be overshooting the target for the majority of researchers and laboratories involved in next generation sequencing analysis – in terms of production capacity and price.

## Conclusions

In conclusion, the Genome Analysis ToolKit 4 offers unprecedented possibilities for parallelization, as shown here by the high efficacy and performance on local cores. It was shown to be several times faster than previous GATK3.8 multithreading with the same multi-core, single CPU, configuration. The complexity of the toolkit – with regard to variant calling – is lowered compared to earlier variant calling with GATK3 Unified Genotyper, as indel realignment is omitted from the latest major version. In addition, GATK4 offers new tools which combines several earlier commands as *pipeline* versions, although still not as stable releases. It comes with “boxed” Spark capabilities *per se* without the need for clusterization in order to directly benefit from the innovation brought by the Broad Institute without Apache Spark installation.

## Declarations

### Ethics approval and consent to participate

Ethics approval is not required according to National Committee on Health Research Ethics in Denmark (http://en.nvk.dk/how-to-notify/what-to-notify) in the current form of methods testing and quality control without any identifiable information.

### Consent for publication

Not applicable.

### Availability of data and material

The panel and amplicon sequencing data used in this study is not publicly shared as it is based on routine clinical hospital tests of leukemia patients with no other purpose in this benchmark than to gather performance metrics. All rights reserved to chief physician, phd, Hans B. Ommen (co-author) and Aarhus University Hospital, Central Denmark Region. Data access can be granted on request if relevant and in accordance to national ethical guidelines. Contact corresponding author for more information. Exome data can be downloaded from The International Genome Sample Resource (www.internationalgenome.org, ID: HG00121)

## Authors’ contributions

ATS and HBO provided clinical sequencing samples for the analysis, MCH performed all analyses described in the paper and wrote the draft. CGN conducted scientific supervision and assisted in finalizing the manuscript. All authors contributed to the manuscript.

## Acknowledgements

The authors would like to thank Dina Mohyeldeen, MD, for a critical view on the final draft. We also wish to thank professor Peter Hokland, Aarhus University Hospital, for continuous scientific supervision and support.

